# Towards designing globular antimicrobial peptide mimics: role of polar functional groups in biomimetic ternary antimicrobial polymers

**DOI:** 10.1101/2020.10.24.353243

**Authors:** Garima Rani, Kenichi Kuroda, Satyavani Vemparala

**Author notes:** Author for communication (Garima Rani).

## Abstract

Using atomistic molecular dynamics simulations, we study the interaction of ternary methacrylate polymers, composed of charged cationic, hydrophobic and neutral polar groups, with model bacterial membrane. Our simulation data shows that the random ternary polymers can penetrate deep into the membrane interior and partitioning of even a single polymer has a pronounced effect on the membrane structure. Lipid reorganization, on polymer binding, shows a strong affinity of the ternary polymer for anionic POPG lipids and the same is compared with the control case of binary polymers (only cationic and hydrophobic groups). While binary polymers exhibit strong propensity of acquired amphiphilic conformations upon membrane insertion, our results strongly suggest that such amphiphilic conformations are absent in the case of random ternary polymers. The ternary polymers adopt a more folded conformation, staying aligned in the direction of the membrane normal and subsequently penetrating deeper into the membrane interior suggesting a novel membrane partitioning mechanism without amphiphilic conformations. Finally, we also examine the interactions of ternary polymer aggregates with model bacterial membranes, which show that replacing some of the hydrophobic groups by polar groups leads to weakly held ternary aggregates enabling them to undergo rapid partitioning and insertion into membrane interior. Our work thus underscores the role of inclusion of polar groups into the framework of traditional binary biomimetic antimicrobial polymers and suggests different mode of partitioning into bacterial membranes, mimicking antimicrobial mechanism of globular antimicrobial peptides like Defensin.

## 1 Introduction

The emergence of antibiotic resistant bacteria presents one of the most daunting challenges faced by scientific community and development of new antibacterial agents is urgently needed in order to fight the resistance mechanisms in pathogenic bacteria ^1,2^. Host defensive antimicrobial peptides (AMPs) are components of the innate immune system of eukaryotes which can effectively kill infecting bacteria, without harming the host cells and without inducing significant resistance ^3^. Such AMPs have been extensively investigated ^4–6^ and indeed, interactions of a variety of AMPs with model cell membranes have been well studied ^7–9^. However, despite their broad spectrum activity and optimised structure, most AMPs are not ideal drug candidates and have many limitations for clinical use ^6^. Accordingly much effort in recent years has been expended into designing synthetic polymers (AMPoly) which can act as effective therapeutic substitutes to the more expensive naturally occuring AMPs. ^10–13^.

Polymer interactions with biological membranes has been intensively studied in recent years ^13,14^, especially to understand their applicability in biomedical and environmental settings. From the perspective of their biomedical relevance, previous studies have shown that the polymers with most optimal antimicrobial activity exhibit weak aggregation behaviour in solution phase ^15–17^. Such weak aggregates allow the individual polymers from the aggregate to insert themselves into a membrane environment facilitating partitioning and disruption of the cytoplasmic membrane to eventually cause cell death. Hence, while aggregation in solution phase is needed to increase the local charge concentration for effective detection of oppositely charged bacterial membranes, the aggregate has to be weak for effective partitioning into the membrane, a crucial initial step in the antimicrobial mechanism. This suggests that the polymers should not have very high hydrophobic content in them, which would promote strongly held aggregates in the solution phase. Given that the antimicrobial polymers are usually of a certain minimum length and the net charge in such systems in typically around +6^18^, from various studies of naturally occurring effective antimicrobial peptides, one can ask the question about how to modulate the hydrophobic content in design of biomimetic AM polymers. Both experimental and simulation studies have shown that modulating hydrophobic content, even by changing the size and length of side chains, can have profound effects on binding and subsequent partitioning into bacterial bilayers suggesting a need to optimize the hydrophobic content of the antimicrobial agents for effective action ^19–22^.

Most of the previous studies on polymers focused on inclusion of only cationic and hydrophobic moieties as the constituents of such designed biomimetic polymers, these two functionalities being considered the minimum requirements for bactericidal activity ^23–26^. However, it is not easy to simultaneously optimize the constituent functional groups to ensure high levels of anti-bacterial activity as well as low hemolytic activity, underlining the importance of exploring polymers having components apart from cationic and hydrophobic groups ^18,27,28^. One possible approach is to introduce additional functional groups into the polymer design to balance the possibility of excessive hydrophobic content. In this regard, there have been recent studies probing the aggregation of ternary polymers in solution suggesting that introduction of polar groups can induce increased solubility and low hemolytic activity ^28^. Furthermore, naturally occurring antimicrobial peptides usually have a distribution of several functional groups in their sequence, including hydrophobic, charged cationic and neutral polar groups ^29,30^. More recent works have started incorporating polar groups, in addition to cationic and hydrophobic groups, in designing antimicrobial polymers ^28,31,32^. For instance, tt has been shown, using ternary nylon-3 copolymers, that replacing hydrophobic or cationic groups or both by hydroxyl residues can result in significantly reduced hemolytic activity as compared to their binary counterparts with only hydrophobic and cationic subunits ^32^. More recently, Mortazavian et al. ^28^ highlighted the active role played by polar groups in reducing the formation of domains of strong hydrophobic monomers in methacrylate random polymers.

Another aspect of interest is the conformational dynamics of the partitioned polymers into the bacterial membrane, which can determine the mode of their antimicrobial mechanism. Literature is available, on both AM peptides and AM polymers, which suggests that either built-in or acquired facial amphiphilicity, in which there is facial separation of charged and hydrophobic groups along the polymer backbone, is one of the hallmarks of effective antimicrobial mechanism ^4,33,34^. These AM peptides are typically helical in nature ^4^, which allow for easy facial segregation of functional groups along the backbone. However, there are other families of antimicrobial peptides like globular defensins ^35,36^, helical AP3^37^ which do not exhibit facially amphiphilic structures but nevertheless have potent antimicrobial activities. The biomimetic AM polymers, largely studied so far and which are composed of only cationic and hydrophobic functional groups, either are designed to have an in-built amphiphilic conformation (usually rigid backbone) or have been shown to have the ability to acquire a facially amphiphilic structure when partitioned into the membrane (usually flexible backbone) ^34,38,39^. However, with the inclusion of new functional groups in the polymer design, one can speculate that facial amphiphilicity may not be possible but can in fact open up new paradigms of designing biomimetic AM polymers with requisite selectivity and antimicrobial action and can be a platform to understand non-facially amphiphilic families of AM peptides mentioned above. It has been suggested that the presence of additional neutral hydrophilic groups likely preclude the formation of amphiphilic conformations in membrane phase ^28^, which hints at a more nuanced polymer-membrane interaction in this case. Nonetheless, there are no experimental or simulation based studies which elucidate the insertion mode of the ternary polymers into bacterial membrane.

In this work, we study the interaction of ternary polymers, composed of charged cationic, hydrophobic and neutral polar groups, with model bacterial membrane using detailed atomistic molecular dynamics simulations and compare them with the action of binary polymers, composed only of charged cationic and hydrophobic groups, on model bacterial membranes. We also investigate the role played by sequence of polar groups in influencing the interaction of such polymers on the bacterial membrane, by considering polymer models having block as well as random arrangements of these groups along their backbones. We find that random ternary polymers penetrate deep into the membrane interior, with even a single polymer markedly affecting the structure of the membrane. Our study also elucidates the strong affinity of the ternary polymer for POPG lipids in membrane phase, resulting in substantial lipid reorganization in polymer neighbourhood in this case. Membrane mode of insertion were also shown to be substantially different for binary and ternary copolymers. Traditional binary copolymer exhibit acquired amphiphilic conformation upon membrane insertion, with cationic ammonium groups localizing close to the interfacial lipid head groups and the hydrophobic moieties buried deeper within the hydrophobic tails of the bilayer. On the other hand, we show that a clear segregation of groups is absent in random ternary polymer case, as had been surmised in previous experimental work ^28^, with the polymer in this case assuming a more folded conformation, staying aligned in the direction of the membrane normal. Finally, we study interactions of aggregates of ternary and binary polymers with bacterial membrane, showing that the weaker ternary aggregate undergoes fairly rapid partitioning and subsequently, membrane insertion while polymer partitioning is robustly obstructed in case of the strong binary aggregates.

## 2 Models and methods

### 2.1 System set up

We performed atomistic MD simulations with explicit water and ions on ternary as well as binary biomimetic copolymers and studied their interactions with model bacterial membrane. Ternary methacrylate random copolymers (referred to as “model T”), consisting of cationic ammonium (amino-ethyl methacrylate: AEMA), hydrophobic alkyl (ethyl methacrylate: EMA) and neutral hydroxyl (hydroxyl methacrylate: HEMA) groups as shown in Fig 1, are modelled with degree of polymerization (DP) = 19^28^. The number of monomers per polymer chain is taken as follows: 6 for AEMA group, 8 for EMA group and 5 for HEMA group. This composition of ternary copolymers has been shown to be optimal for antimicrobial activity while also having significantly reduced hemolytic activity ^28^, a major consideration for designing anti-bacterial agents. We also consider a second ternary polymer model (referred to as “model Tb”), which has the same proportion of groups, but is sequentially different from model T polymer. In this case, the AEMA groups and EMA groups are arranged randomly but the HEMA groups are arranged in a block sequence at one end of the polymer backbone. Further, to highlight the role of inclusion of neutral polar functional groups, we also perform control simulations without them, involving only binary compositions of cationic (AEMA) and hydrophobic (EMA) monomer units in random configuration and with same degree of polymerization as ternary polymers. The number of monomers per chain in the binary polymer is taken to be 6 (AEMA) and 13 (EMA). For all the three model polymers, the number of cationic side chain monomers is fixed to be 6 per polymer, in agreement with results of Mortazavian et al ^28^, where it was found that despite different EMA composition, the minimum inhibitory concentration (MIC) value of all polymers starts to level off at ~ 30% of AEMA groups and that the proportion of cationic groups above 30 % does not increase the antimicrobial activity of the polymer. Further, this proportion of cationic group is also comparable to net positive charge of natural AMPs ^40^. The proportion and sequence of the groups along the three model polymers is summarised in Table 1.

**Fig. 1.**
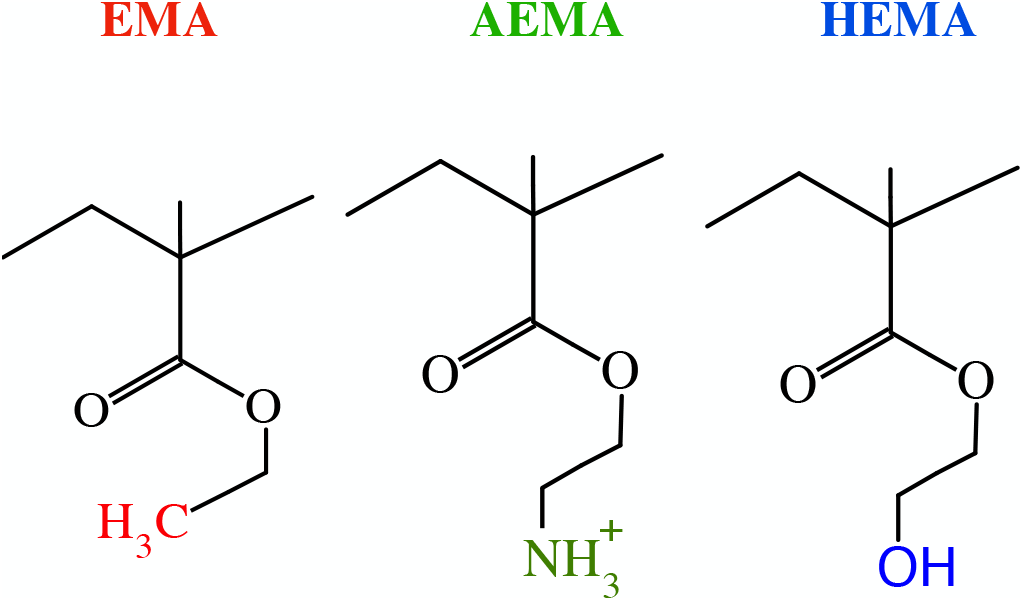
Chemical structures of EMA, AEMA and HEMA groups considered in the model polymers.

**Table 1.** Proportion and sequence of AEMA (A), HEMA (H) and EMA (E) monomers in the polymer models (T,Tb and B). In all the model polymers, degree of polymerization (DP) = 19 and the number of cationic side chain groups are fixed to be 6 per polymer. Model T and Tb are compositionally same, but differ in sequence of the three groups along the polymer backbone

| Model copolymers | DP | Group proportion (AEMA, HEMA, EMA) | Group Sequence |
| --- | --- | --- | --- |
| Ternary (T) | 19 | +6, 5, 8 | E-H-A-E-H-A-H-E-A-E-E-A-E-H-H-A-E-E-A |
| Ternary (Tb) | 19 | +6, 5, 8 | E-A-A-E-E-A-E-E-A-E-E-A-E-A-H-H-H-H-H |
| Binary (B) | 19 | +6, 0, 13 | E-E-A-E-E-A-E-E-A-E-E-A-E-E-E-A-E-E-A |

For bacterial membrane, the starting configuration was taken from a pre-equilibrated membrane patch consisting of 38 POPG and 90 POPE lipid molecules per leaflet, in similar ratio (3:7 POPG to POPE) as the inner membrane of the Gram negative bacteria ^41^, constructed using CHARMM-GUI’s Membrane Builder module ^42^ and used in our previous simulations ^17,43^.

### 2.2 Simulation protocols

Single polymers were simulated in a box (≈ 50 Å × 50 Å × 50 Å) of TIP3P water model before placing them near the membrane environment. Since the total charge of a single polymer in each of the system was +6*e*, an appropriate amount of NaCl salt was added to neutralize the systems and maintain 150 mM salt concentration to mimic physiological conditions in each case. The total number of atoms in this simulation system was ~ 10500. All simulations were performed with the NAMD 2.9 simulation package ^45^. Each polymer system was first energy minimized for 1000 steps with the conjugate gradient method and simulations were then performed with 2 fs timestep in the isothermal-isobaric (NPT) ensemble for 200 ns.

#### Single polymer-membrane systems

To construct the membrane-polymer system, a randomly chosen equilibrium configuration, from above solvent simulations, of each of the polymer (models T, Tb and B) was placed in the vicinity of one of the bilayer (referred to as upper leaflet) along the membrane normal in the TIP3P water phase and required number of counterions of Na^+^ and Cl^−^ were added to neutralize the system and maintain 150 mM salt concentration. These three different polymer-membrane ensembles (consisting of membrane and polymer models T, Tb and B, respectively) were simulated with periodic boundary conditions in the isothermal - isobaric (NPT) ensemble. The latest CHARMM force fields CHARMM 36 were used for lipid molecules ^46^. The parameter values for the polymers were adopted from the CHARMM force field ^47^ and previous simulations ^17,20,48^. The total number of atoms for the polymer-membrane simulations was ~ 71000. Constant temperature was maintained at 310 K, which is above the main-phase transition temperature of both POPE and POPG lipid molecules ^49^ and a pressure of 1 atm was maintained through Langevin piston ^50,51^. Electrostatic interactions were calculated by the Particle Mesh Ewald method ^52^ and the cut-off for non-bonded interactions was set to 12 Å, with smoothing starting from 10 Å. The polymer-membrane systems were first energy minimized using conjugate gradient method and simulations were conducted with a 2 fs timestep. Polymer in each case was initially subjected to harmonic restraint which was gradually reduced to zero over 2 ns. System with random ternary model T-membrane system was simulated for 900 ns, whereas the ternary Tb-membrane and binary B-membrane systems were simulated for 700 ns. Simulation details for the single polymer-membrane systems are summarised in Table 2.

**Table 2.**
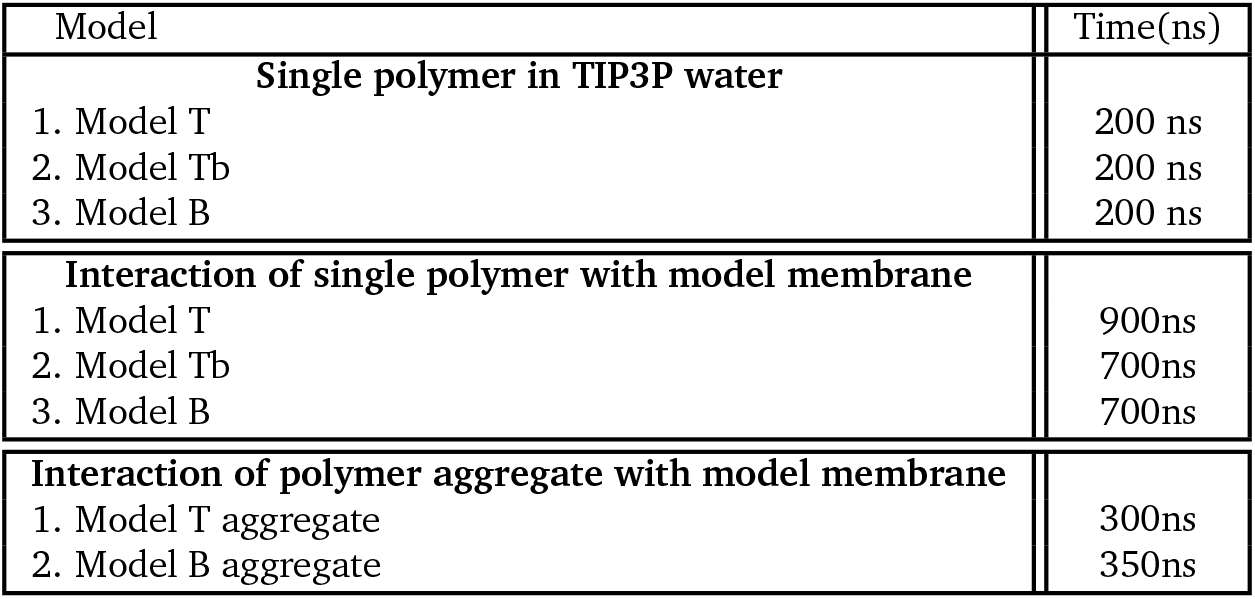
Simulation times in each case for the single polymer-water solution, single polymer-membrane and polymer aggregate-membrane systems. The number of POPG and POPE lipids per leaflet were 38 and 90 respectively.

| Model | Time(ns) |
| --- | --- |
| **Single polymer in TIP3P water** |  |
| 1. Model T | 200 ns |
| 2. Model Tb | 200 ns |
| 3. Model B | 200 ns |
| **Interaction of single polymer with model membrane** |  |
| 1. Model T | 900ns |
| 2. Model Tb | 700ns |
| 3. Model B | 700ns |
| **Interaction of polymer aggregate with model membrane** |  |
| 1. Model T aggregate | 300ns |
| 2. Model B aggregate | 350ns |

#### Polymer aggregate-membrane systems

To study membrane interactions of polymer aggregates, multiple ternary polymers (model T) and binary polymers (model B) were first dispersed in a box of water and sufficient counterions to maintain 150 mM salt concentration. This system was simulated for 150 ns in NPT conditions, from which we then extracted the largest formed stable aggregates of binary and ternary polymers. Both these aggregates consisted of four polymers, four ternary polymers (denoted T1, T2, T3, T4) in case of the ternary model T aggregate and four binary polymers (denoted B1, B2, B3, B4), in case of the binary model B aggregate. These aggregates of ternary and binary random polymers were then placed in the vicinity of the upper leaflet and simulated for atleast 300 ns in NPT ensemble. The total number of atoms for the aggregate-membrane simulations was ~ 80000. All the parameters and conditions for the aggregate-membrane systems were kept the same as the single polymer-membrane systems. Simulation details for the polymer aggregate-membrane systems are summarised in Table 2.

For analysis and visualization, the VMD package ^53^ analysis tools was used. All the analysis for the membrane-polymer systems is performed using plugins and TCL scripting language embedded with VMD.

## 3 Results

### 3.1 Polymer insertion modes in membrane phase

Both ternary and binary polymers, owing to the presence of amino groups which are cationic in nature, get attracted towards PG lipid groups of the bacterial membrane through electrostatic interactions ^40^. In our case, for all three polymer-membrane systems that were simulated, single polymer placed in water quickly approached the membrane patch within a few nanoseconds and localized close to the nearest membrane leaflet (referred to as the upper leaflet). However, after contact with the leaflet, significant differences were observed in the conformation of the three polymers, which we delineate in this section. Our objective is to study the role of the presence of the HEMA groups and their arrangement along the polymer backbone in governing the polymer-membrane interaction.

In Fig. 2, the time evolution snapshots of the polymer-membrane systems is depicted over the entire simulation timescale. It is clear that different polymer models display remarkably different membrane interaction mechanisms in terms of the conformation they achieve during membrane partitioning. For the random ternary polymer (model T with AEMA, EMA and HEMA groups forming random ternary copolymer), though the polymer comes into contact with the membrane surface within first few nanoseconds, it can be seen that the complete partitioning of the polymer into the membrane interior is hampered. This is due to the competing interactions between the polymer function groups and various groups of the POPE-POPG membrane system. The presence of both AEMA and HEMA groups (and consequently lowered hydrophobic EMA content) most likely encourage the polymers to be present largely at the membrane-water interface as both the charged AEMA and polar HEMA groups can have favourably interactions with the water and the head-group region of the membrane. The interaction polymer T samples different conformations and settles into a globular structure (Fig. 2A) at around ~ 350 ns, with the polymer penetrating the membrane from both of its ends, gripping the membrane akin to a holdfast. Subsequently, the polymer assumes an even more folded conformation in the membrane interior (Fig. 2A), with the EMA groups near both the inserted ends of the polymer coming in contact with each other, while projecting towards the membrane core due to favourable hydrophobic interactions.

**Fig. 2.**
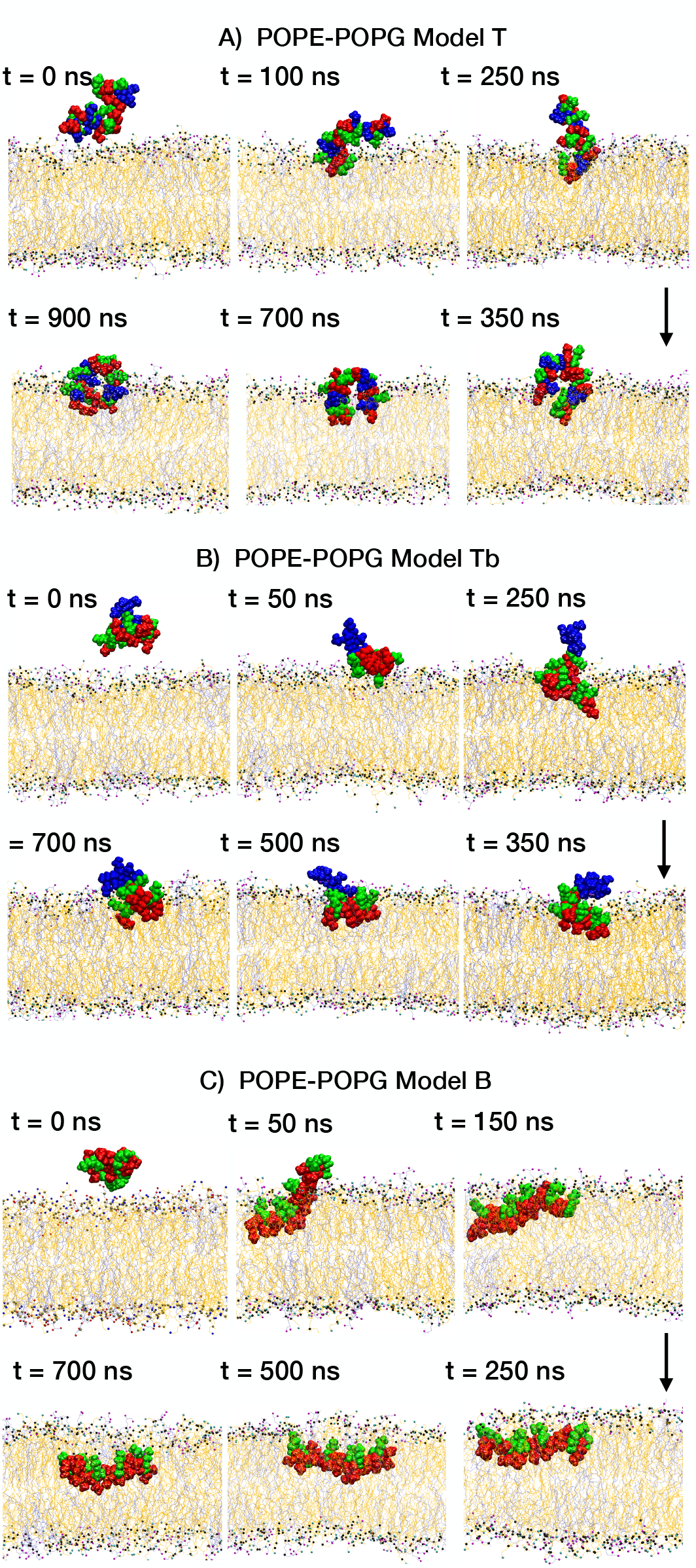
Representative snapshots of A) model T, B) model Tb and C) model B, interacting with the bacterial bilayer. The POPE lipids are coloured orange and POPG lipids are shown in ice-blue colour; the lipid head groups are oxygen (magenta), nitrogen (cyan) and Phosphate (black). The cationic, hydrophobic and polar groups of the model polymers are shown in green, red and blue colour respectively.

To gain insight into the role played by sequence of polar functional groups in interactions with bacterial membrane, we also performed simulations involving ternary polymer model Tb, having random arrangements of EMA and AEMA groups, and a block of HEMA groups clustered at one end (Table 1). In contrast to the model T polymers, the clustering of the polar HEMA groups to one end of the polymer results in a very different conformational sampling of the polymers. While the randomly distributed charged AEMA and hydrophobic EMA groups try to adopt a facially amphiphilic structures, the clustered polar HEMA groups predominantly stay in the solution phase (Fig. 2B), clearly due to favourable polar interactions with water.

To contrast the results for the ternary polymer systems and in particular to understand the role of HEMA groups, a control simulation is performed taking the same length of the polymer but with only binary composition of charged AEMA and hydrophobic EMA groups (polymer B). As in the previous two cases, the presence of cationic charged groups ensures the contact of the polymer B with the membrane surface. In a complete departure with the polymers T and Tb, the binary polymer B partitions completely into the POPE-POPG membrane and adopts a near perfect facially amphiphilic structure and resides in the membrane core, parallel to the membrane surface. In this conformation, the hydrophobic groups of the polymer B are projected towards the membrane core and the cationic ammonium groups projected (or snorkel up) towards the water - membrane interface (Fig. 2C). This adoption of facially amphiphilic structures in polymers with binary composition excellently mimics our previous observations of different polymer length chains and various side chain spacer groups ^17,20,54^, although it is to be noted that in our previous studies, polymers with ethyl cationic side arms did not display a tendency to form facially amphiphilic conformations, suggesting that both monomer composition as well as polymer length can affect the adoption of such conformations in membrane phase. Further, due to higher hydrophobic content of the polymers studied here, in addition to longer simulation time scales, the penetration depth of the partitioned polymer B inside the POPE-POPG is significantly higher than our previous results.

These results clearly suggests that for the polymer conformations, not only the presence of various functional groups is important, their sequence along the polymer backbone is also important. Since the antimicrobial mechanism of these biomimetic polymers may depend crucially on the polymer conformations, insights obtained here about the composition and sequence of the functional groups can aid our understanding and design of the AM polymers. Our aim now is: (a) to quantify the strength of the polymer-membrane interactions for the ternary polymer models and compare it to the binary model, (b) to study the polymer conformation and its evolution in membrane environment and to compare it to their conformation in solution phase and (c) to assess the effect, if any, that the polymer has on the structural properties of the membrane, particularly in the ternary case.

### 3.2 Conformations of membrane partitioned polymers

In this section, we probe in detail the polymer conformation and its evolution, as it interacts with the membrane. For this, we first map the spatial distribution of the various constituent moeities of the polymer models by computing their density profiles along the membrane normal (here z-axis). The z-density profiles averaged over last 50 ns for the three model polymer-membrane systems are shown in Fig. 3. For the model T polymer, the presence of hydroxyl side chain groups in a random configuration obstruct a clear division of the groups and significant overlap between the peaks corresponding AEMA, EMA and HEMA groups is observed in the z-density profile as shown in Fig. 3A. On the other hand, the z-density plot for model Tb-membrane system show a much lower overlap between AEMA and EMA groups, while the HEMA moeities remain outside the membrane interior (Fig. 3B), consistent with observations in the previous section. Thus, block arrangement of the HEMA groups clearly facilitates spatial segregation of groups, with the EMA groups preferring the membrane interior, the AEMA groups staying closer to the lipid head groups and the polar HEMA groups preferring aqueous phase. We note that model B polymer is inserted into the bacterial membrane with well separated and least overlapping average peaks between AEMA and EMA groups, indicating the spatial segregation of the two groups (Fig. 3C). Indeed, previous simulation studies have demonstrated that binary AM polymers can adopt facial amphiphilicity upon interacting with cell membranes, even though the polymers may themselves lack a secondary structural conformation ^17,20,54,55^. We also note that the cationic groups align well with the phosphate head groups of the lipids and the hydroxyl group of POPG lipid, highlighting their preference for interactions with the charged head groups of the bilayer. The plots in Fig. 3 also show the depth at which the polymers are anchored within the membrane leaflet. Density data shows peak position of cationic AEMA group is deeper for model B and model Tb polymers as compared to the model T case, although further comparing the two ternary models reveals that the three groups (EMA, AEMA, HEMA) of the model T polymer have more average contacts inside the membrane, compared to model Tb polymer, since block HEMA moeities remain localized outside the membrane.

**Fig. 3.**
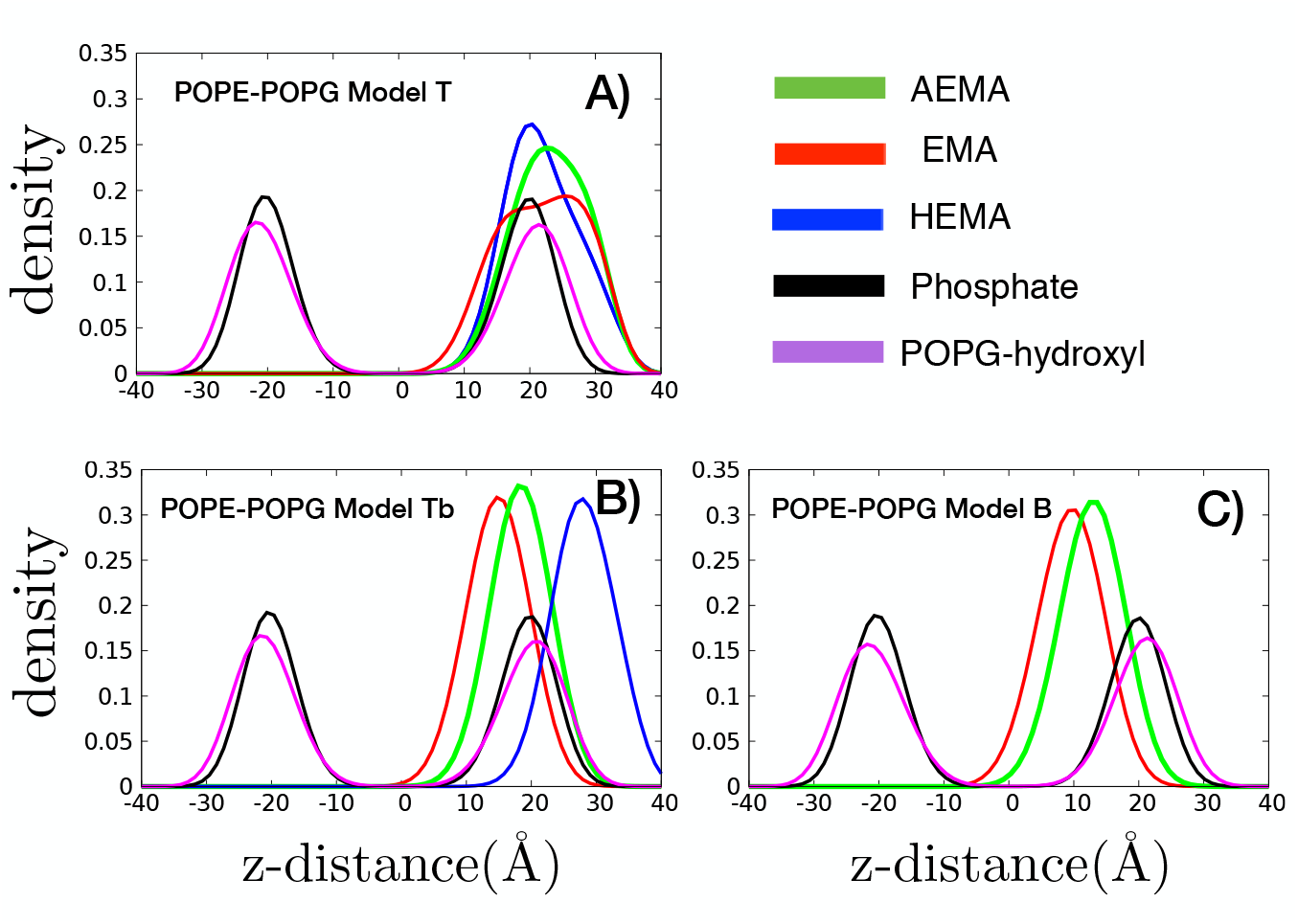
Density profiles of various components of lipid-polymer systems, A) POPE-POPG model T, B) POPE-POPG model Tb and C) POPE-POPG model B, averaged over last 50 ns of MD simulations. The green line represents the polymer charged group, red colour represents the polymer hydrophobic group and the blue colour marks the polymer polar group. The density profile of phosphate atoms, hydroxyl groups of POPG lipids are are shown in black and magenta, respectively.

To characterize the onset of the facial amphiphilicity and the placement of the different polymer functional groups with respect to lipid head groups along the membrane normal, we monitor the time evolution of the center of mass of the most partitioned groups of the AEMA and EMA groups of polymer models T, Tb and B and HEMA group of models T and Tb polymer, with respect to the bilayer in Fig. 4 (note that for the model T polymer, the two most partitioned HEMA residues associated to the two ends of the polymer are plotted). We also compute the evolution of the center of mass of all the groups in the three model polymers with respect to the membrane lipid phosphate head groups in Fig.1 of the Supplementary Information. We observe that in all the three cases, polymer placed in water phase quickly approached the membrane-water interface. This is due to the presence of similar proportions of cationic AEMA groups in all the three polymer models, which get attracted towards the anionic head groups of POPG lipids. Interestingly, the hydrophobic side chain and the cationic groups undergo a clear flip just upon entering the upper leaflet surface. This is in good agreement with the flipping observed in previous work ^17^. As noted above, a clear segregation of groups is absent for the model T polymer (Fig. 3A). The positional inclination of the HEMA groups is understood by examining the evolution of the center of mass for groups in the model Tb polymer, where these remain outside the membrane interior throughout the simulation time scale, showing a preference for interactions with water. The time evolution of the cationic and the hydrophobic groups in this case show a flipping of the EMA and AEMA groups just upon membrane insertion following which the EMA groups consistently stay below the AEMA groups (Fig. 3B). This is similar to the control case of binary polymer model B, in which time evolution of the center of mass well displays the adopted amphiphilic conformation for model B polymer, this facial amphiphilicity being initiated once the polymer inserts into the membrane interior and persisting throughout the simulation time after this (Fig. 3C). However, model Tb moeities maintain a shallower depth in the membrane as compared to the model B case, because of the presence of clustered HEMA groups in the polymer.

**Fig. 4.**
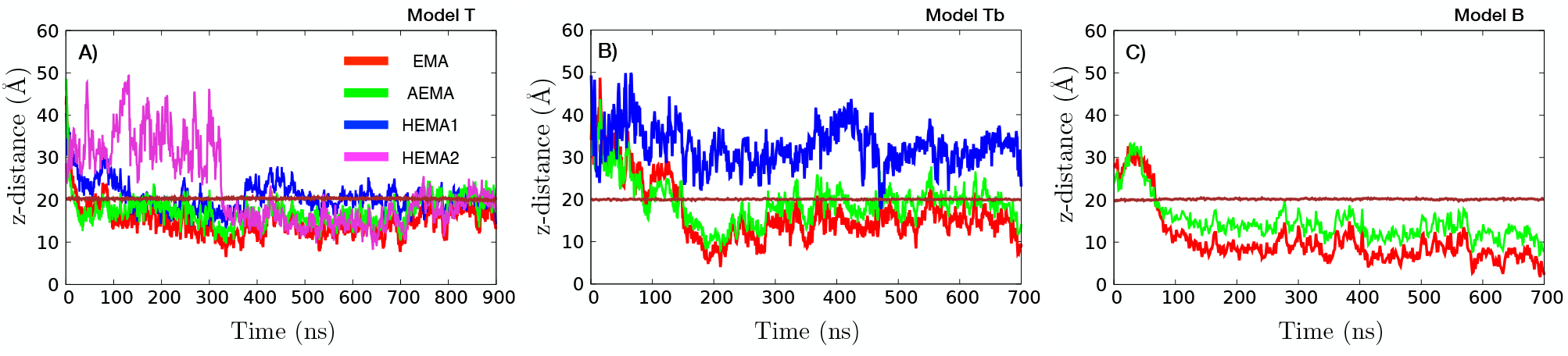
Time evolution of center of mass (z-component) of the hydrophobic EMA, cationic AEMA and hydrophilic HEMA groups that are lowermost at the end of simulation time. Note that for model T polymer, we plot the center of mass of two HEMA groups, denoted HEMA1 and HEMA2, placed at the two ends of the polymer. The HEMA2 group is placed at the end of the polymer which inserted into membrane at around t ~ 350 ns.

#### 3.2.1 Morphological changes of the polymers in solution and membrane environment

The differences in the conformations of random polymers (models T and B) in solution phase and in lipid phase are compared to understand the effect of environment on the morphology of the polymers. The distribution and time evolution of the radius of gyration (*R_g_*) values for models T and B polymers are shown in Fig. 5. The observed lower values of *R_g_* for binary polymer in solution phase, as compared to ternary polymer (Fig. 5A) can be attributed to the higher hydrophobic content in the model B polymer. Accordingly, the model T polymer due to presence of higher hydrophilic content (both charged AEMA and polar HEMA functional groups) adopts more extended conformations in the solution (Fig. 6A (top right)). However, when the polymers are near the water-membrane or partitioned into the membrane environment, the conformations of binary and ternary polymers are significantly different. While the ternary polymers, with polar HEMA groups replacing some of the hydrophobic groups, adopt a predominantly globular structure at the water-membrane interface (Fig. 6A (bottom right), the binary polymers partition completely into the POPE-POPG membrane and take strikingly facially amphiphilic and extended structure (Fig. 6B (bottom right), as is also seen in the single peak at higher *R_g_* values (Fig. 5C). The random ternary polymer (model T), shows two peaks in values of *R_g_* (Fig. 5A) and the time evolution depicts that the *R_g_* values initially stayed quite similar to the values of the model B polymer before undergoing a significant fall at ~ 350 ns and then continuing a downward trend. This is due to its initial extended conformation, having penetrated the membrane from one end while staying perpendicular to the membrane surface, before the polymer folds itself to enter the membrane with its other end at ~ 350 ns and gradually assuming increasingly folded conformations (Fig. 2A).

**Fig. 5.**
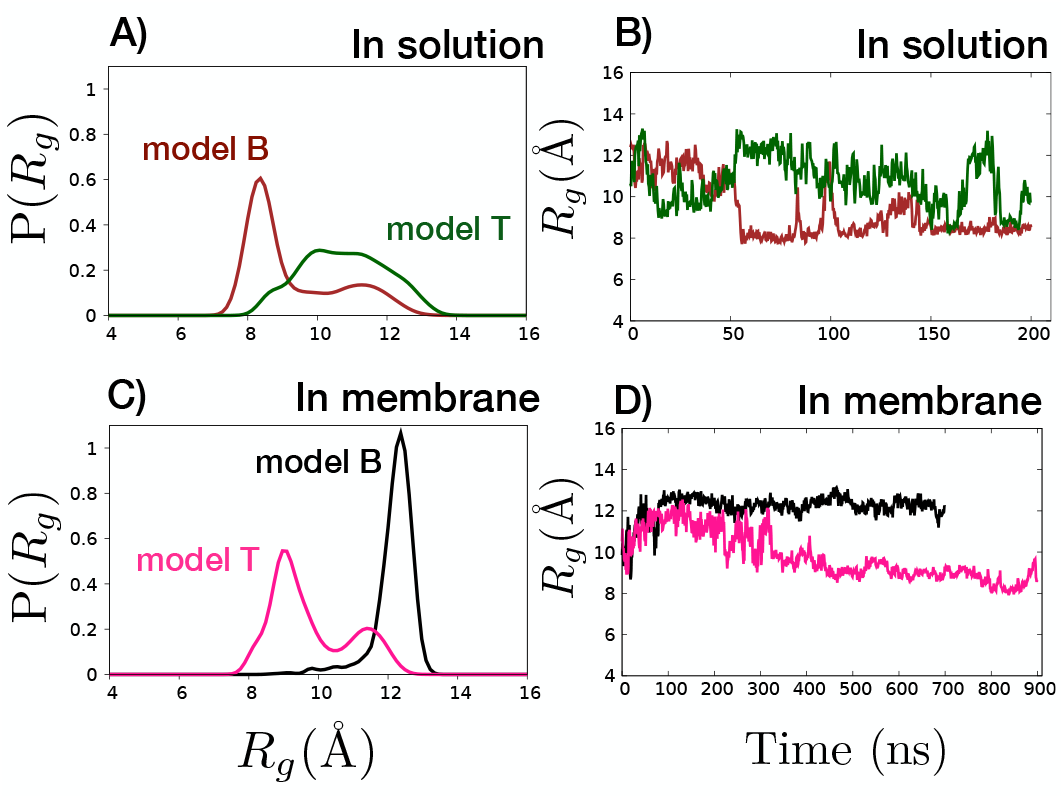
A) Distribution of radius of gyration (*R_g_*), with data sampled over the entire trajectory, for polymer models T and B is shown in solution phase. B) Time evolution of *R_g_* is shown for both the polymers in solution phase. C) Distribution of radius of gyration (*R_g_*) for polymer models T and B is shown in membrane phase. D) Time evolution of *R_g_* is shown for both the polymers in membrane phase.

**Fig. 6.**
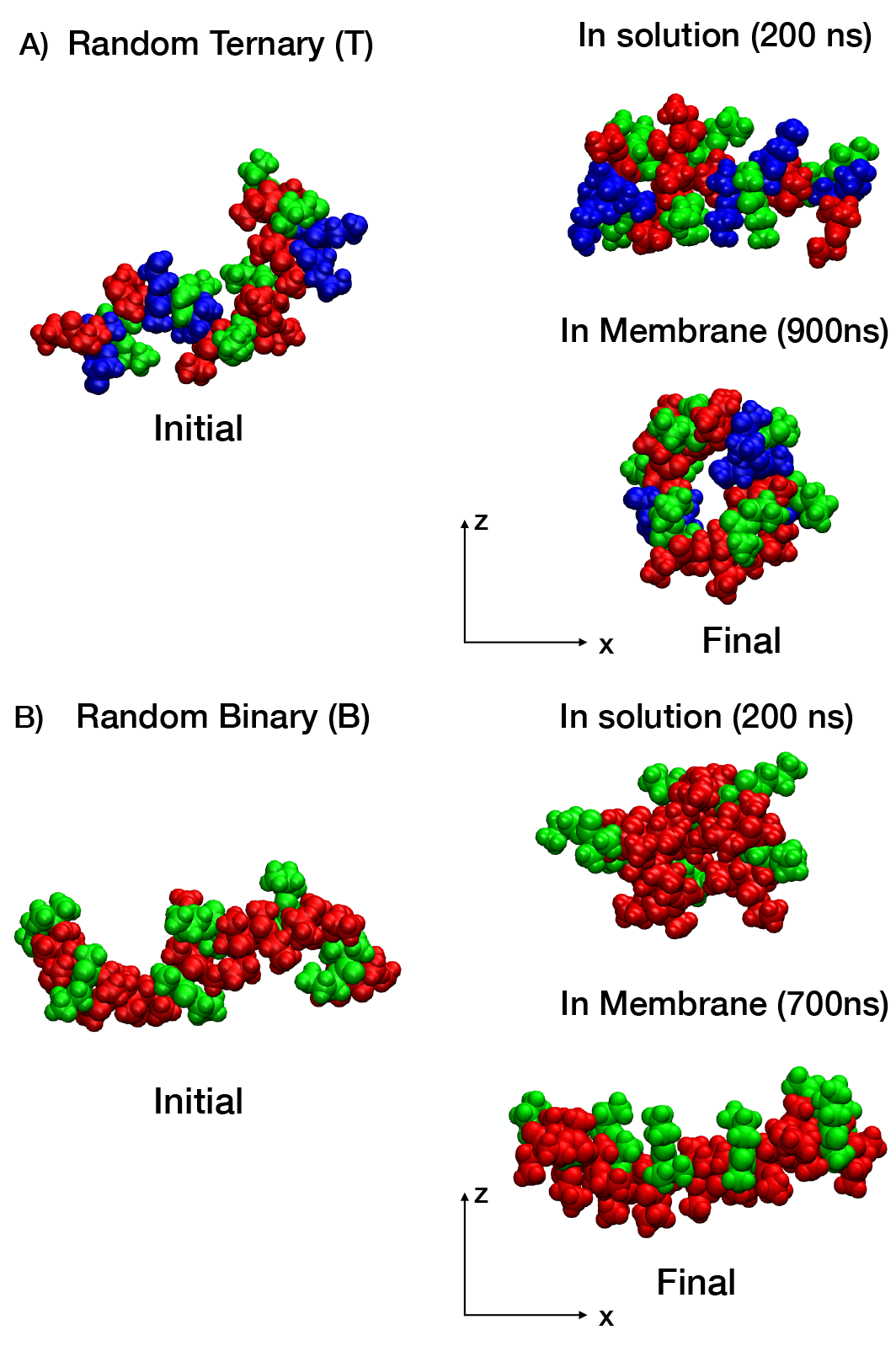
A) Snapshots of initial (left) and final (right) conformations of ternary polymer model T in solution (top right) and in bacterial membrane (bottom right) environments. B) Snapshots of initial (left) and final (right) conformations of binary polymer model B in solution (top right) and in bacterial membrane (bottom right) environments. Cationic, hydrophobic and neutral polar groups are colored green, red and blue, respectively.

### 3.3 Polymer membrane interactions

As noted in the previous section, the replacement of some of the hydrophobic groups in the polymer B by polar HEMA groups to obtain either model T or model Tb results in complete or partial loss of acquired facial amphiphilicity and strikingly different conformations of the partitioned polymers. In particular, the effect of sequence of presence of polar HEMA groups along the polymer backbone has a strong effect on whether the polymer takes a folded conformation (in model T) or somewhat extended conformation (in model Tb). What is conclusive in either of the two cases is that the addition of polar HEMA groups inhibit deeper penetration of the polymers into the membrane environment and hint strongly at a completely different mode of antimicrobial mechanism than those of simple binary polymers and may not require acquiring of facially amphiphilic conformations.To understand differences in propensities of interactions between functional groups of the polymers and membrane, we perform two sets of analyses: time evolution of contacts between polymer and water/membrane and the radial density distributions *g*(*r*).

For time evolution of environmental fractional contacts, we calculate the fraction of water atoms, POPE and POPG lipid atoms in the vicinity of the model polymers as a function of time. The time series data of the fraction of water and lipids within cutoff of 7 Å around EMA and AEMA groups of the model polymer, is plotted in Fig. 7. Ions are not considered in this analysis, since they form a very small fraction of the total atoms in the system. Fig. 7 highlights the evolution of the POPE and POPG lipid species around the groups of the polymer, as the model polymers bind and insert into the membrane leaflet. The polymers initiate their journey surrounded by water but within a few nanoseconds, water contacts are replaced by contacts with lipids species. However, further probing of these analysis, brings out the remarkable role of the composition and sequence of the functional groups on the polymer-membrane contacts. For model T polymer which adopts the most compact and folded conformation at the water-membrane interface, compared to the other two models, the total contacts between polymer and membrane are somewhat comparable. The individual contact evolution of different functional groups are also similar, owing to the random distribution of functional groups along the polymer. For the model Tb, which has the same composition as that of model T, but with clustering of polar HEMA groups, the contact time evolution is strikingly different from model T Fig. 7 (D, E,F). The hydrophobic EMA groups follow a rapid partitioning into the membrane to reduce the energy costs of presence in solution phase, something that random presence of HEMA groups did not allow in the case of model T polymers. This can be seen from significant decrease of EMA-water contacts and increase in EMA-membrane contacts. Though the contact profile of charged AEMA groups in model Tb with water/membrane is not significantly different from that of model T polymer, the contact profile of polar HEMA groups is strikingly different in model Tb polymer as compared to model T. The HEMA groups maintain the contacts with water through out the simulation time, even when the polymer partially partitions into the membrane, strongly suggesting that polar HEMA groups prefer solution phase rather than membrane phase and this can have profound consequences on the polymer conformation at the membrane interface, as seen already. Finally for the model B polymers, which partition deepest into the membrane core, the EMA-water contacts decrease sharply with a significant increase in EMA-membrane contacts. Even in the case of charged AEMA groups, for model B polymer, there is a significant decrease in AEMA-water contacts as the penetration of the polymer into the membrane core increases with time. It is also to be noted that, remarkably, in case of the model T polymer, POPG contacts consistently dominate POPE contacts for EMA, AEMA and HEMA groups, even though as we noted, the number of POPE lipids is more than twice the POPG lipids in the model bacterial membrane system considered here.

**Fig. 7.**
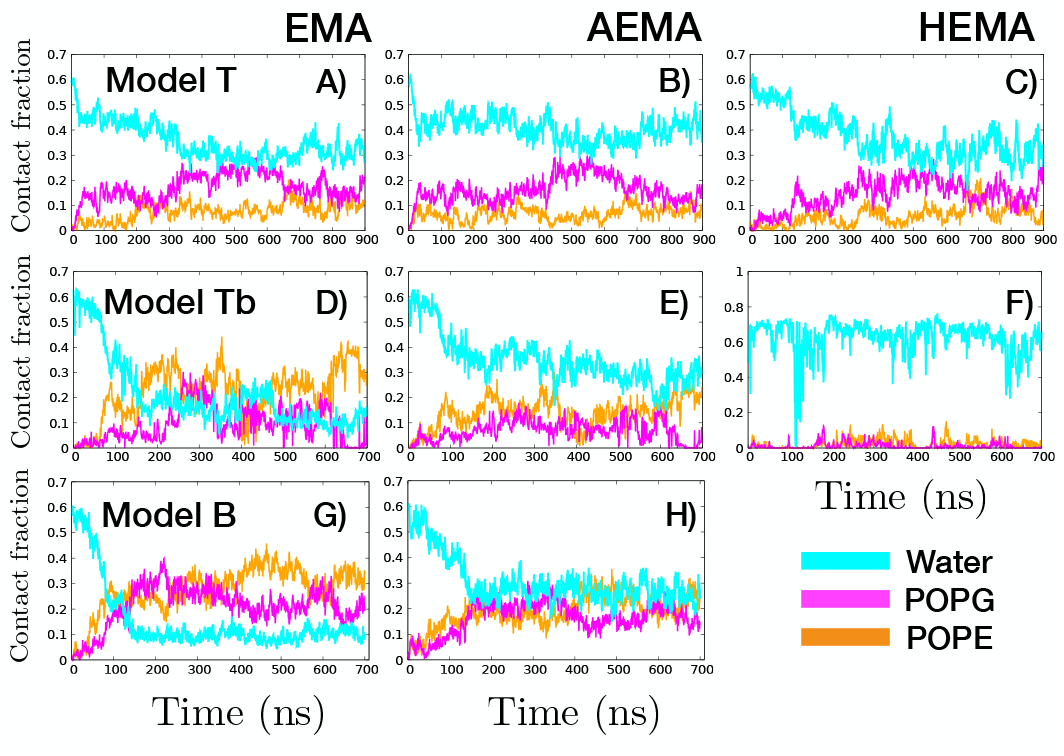
Time evolution of the fraction of contacts that are water and lipids (POPE/POPG) surrounding (A, C, F) EMA, (B, D, G) AEMA and (E, H) HEMA groups, for ternary (T,Tb) and binary (B) polymers. We calculated the fraction of contacts of the polymer that are with water, lipids and polymer particles within cut-off distance of 7Å. Remaining fraction in the figure are contacts of the polymer with itself.

More insight into the interaction between functional groups of polymers and membrane is obtained via *g*(*r*) calculation for the cationic ammonium groups and the hydroxyl groups of the polymers with the headgroup constituents of the POPE-POPG membrane. Fig. 8. Expectedly, this data depicts the strong propensity of the cationic groups (AEMA) of all the three polymer models to bind to overall negatively charged lipid head groups POPG confirming the primary role of charged functional groups of the polymers in identifying the bacterial membranes via attractive electrostatic interactions. However, the interactions of the polar HEMA groups with the negatively charged POPG headgroups crucially depends on their sequence in the polymers: while in model T, where they are distributed randomly, the HEMA groups interact equally strongly with POPG headgroups as their charged AEMA counterparts, in model Tb, where they are clustered, the HEMA-POPG interactions is strikingly suppressed. This is because in the latter case, as seen in the figures Fig. 2B and Fig. 4, the clustered HEMA groups largely prefer the water enviroment and do not partition at all into the membrane phase.

**Fig. 8.**
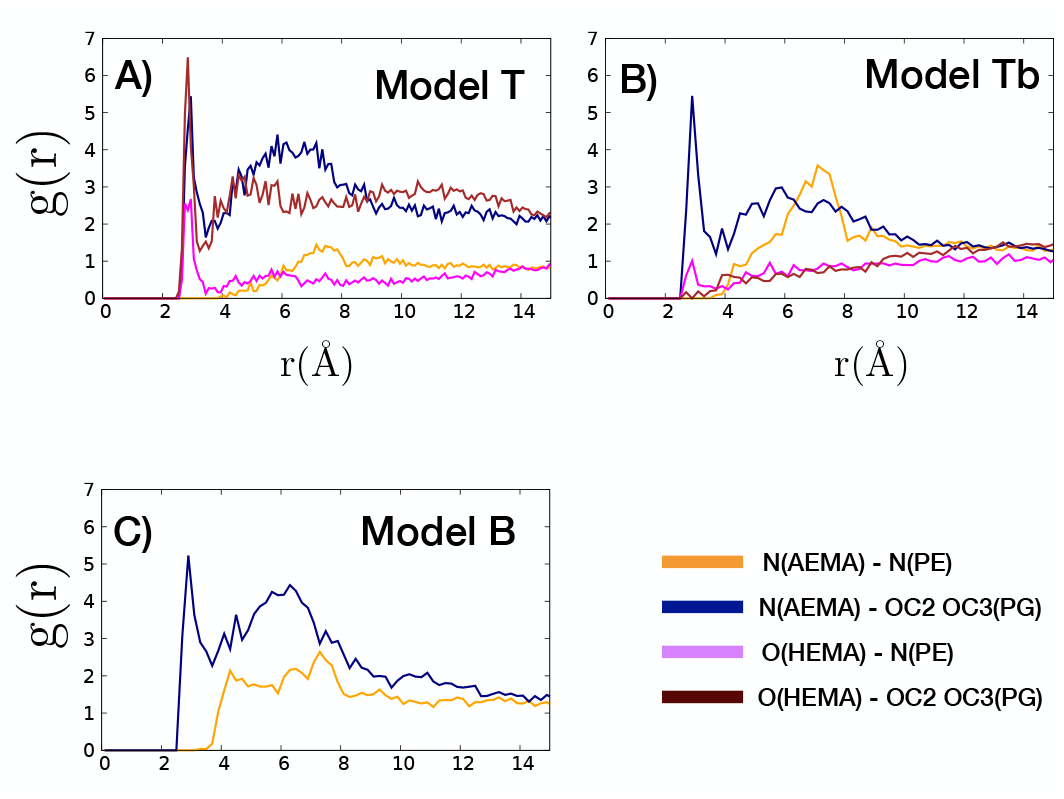
The radial density distribution functions for ammonium and hydroxyl groups of polymer models around the head groups of POPE and POPG lipids. Strong propensity of the cationic groups of the polymers to be around the negative head groups of POPG lipids is shown. Also, HEMA groups of the random ternary polymer prefer interactions with lipid head groups due to their polar nature.

To further understand the role of positioning or sequence of polar HEMA groups on the interactions between the partitioned polymers and the membrane systems, we specifically consider the non-bonded interactions between HEMA groups of the polymers and the bacterial membrane for both models T and Tb as in Fig. 9A and Fig. 9B. We observe that in the model T case, the HEMA group shows higher van der Waals attraction with the bacterial membrane compared to model Tb (block HEMA in ternary copolymer). Even the electrostatic interaction is consistently more attractive in the model T case. From this we can conclude that polymer-membrane interaction is more robust in case of the random ternary polymer, with a block arrangement of polar HEMA groups clearly limiting effective interaction of the model Tb polymer with the bacterial membrane.

**Fig. 9.**
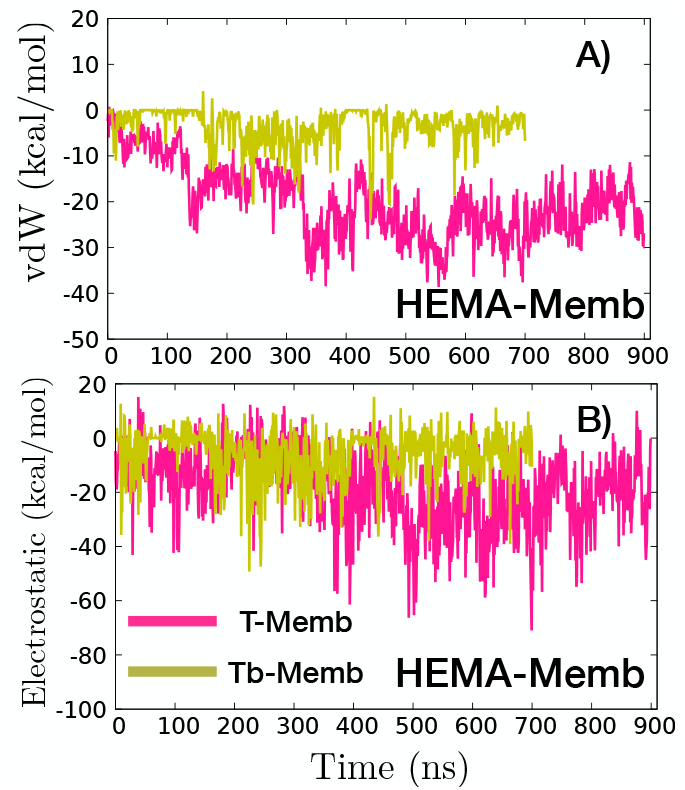
A) Van der Waals and B) electrostatic energy between the HEMA groups and the bacterial membrane for ternary polymer - model T and ternary polymer with block HEMA - model Tb. The van der waals energy is clearly more attractive for polymer model T, where HEMA groups are randomly distributed along the polymer backbone, while in model Tb, the block HEMA groups seem to collectively prefer not to penetrate the bacterial membrane surface and favour water environment.

We can then conclude that the lack of a clear segregation of groups in case of the random ternary polymer is underpinned by two competitive effects involving the HEMA groups, as demonstrated by the energy analysis and the radial density distribution data - preferable interactions with water, which constrain insertion into membrane and attractive interactions of HEMA group with membrane lipids. A block arrangement of the HEMA groups, as in case of model Tb polymer, mitigates this effect by largely favouring interactions with water and thus remaining at and above the membrane-water interface.

### 3.4 Lipid reorganization in presence of polymer

In the previous sections, we have shown that the polymer conformations crucially depend on composition, sequence of the functional groups and the environment (solution or membrane phase). Given the different and preferential interactions of the three different polymer models with POPE-POPG membranes, we next probe whether the polymer interactions have affect on the membrane structure. In previous simulation and experimental works ^17,56–59^, it has been suggested and shown that one of the modes in which biomimetic antimicrobial agents, especially those based on random conformations in solution phase, affect the integrity of the membrane by sequestering/clustering the negatively charged lipids, leading to large scale reorganization and coarsening of the membrane with consequent domain defects, which are exploited by such membrane active AM agents towards final lysis of bacterial cells.

To probe such possible reorganization of the POPE-POPG membrane upon polymer interaction and partitiong, 2D radial density plots (shown in Fig. 10) to measure clustering of POPG lipid molecules is computed for models T and B polymers. The polymers T and B interact with the designated ‘upper’ leaflet of the membrane system. The results show that when ternary polymer interacts with the POPE-POPG membrane, they lead to clustering of the POPG lipids molecules as evidenced by the appearance of the first peak around 4.5 Å, which is absent in the control polymer-free membrane simulation (blue line in Fig. 10 A). It is also to be noted that for the lower leaflet case, where no polymer is present, the 2D radial distribution of POPG molecules remains more or less the same, further underscoring the role of polymer interaction with the membrane on the possible clustering of the POPG molecules. However, for the model B polymer interactions with the POPE-POPG membrane systems, there is no clustering of the POPG molecules is observed in either upper leaflet (where the polymer partitions) and the lower leaflet with and without polymer as seen in Fig. 10 (C,D). This gives a possible insight that due to the preponderance of the hydrophobic EMA groups in model B polymers, the partitioning is primarily driven by the dominant hydrophobic interactions whereas the folded structure and constrained partitioning of the model T polymer recruits the charged POPG molecules around it, significantly reorganizing the membrane in its vicinity. In Fig. 11, a representative snapshot of the POPE (orange color) and POPG (iceblue color) lipids that are within 5 Å around the polymer is shown highlighting the observation that POPG lipids are present in a higher concentration around the polymer as compared to POPE molecules. Thus, our analysis highlights the considerable membrane remodelling that occurs upon insertion and binding of model T random ternary polymer. This is particularly significant since it has been shown that presence of PGs in bacterial membrane significantly increases its stability ^60^, so it is likely that disruption of POPG lipids in the bacterial membrane leads to membrane destabilization, thus promoting a robust antimicrobial action of such polymers.

**Fig. 10.**
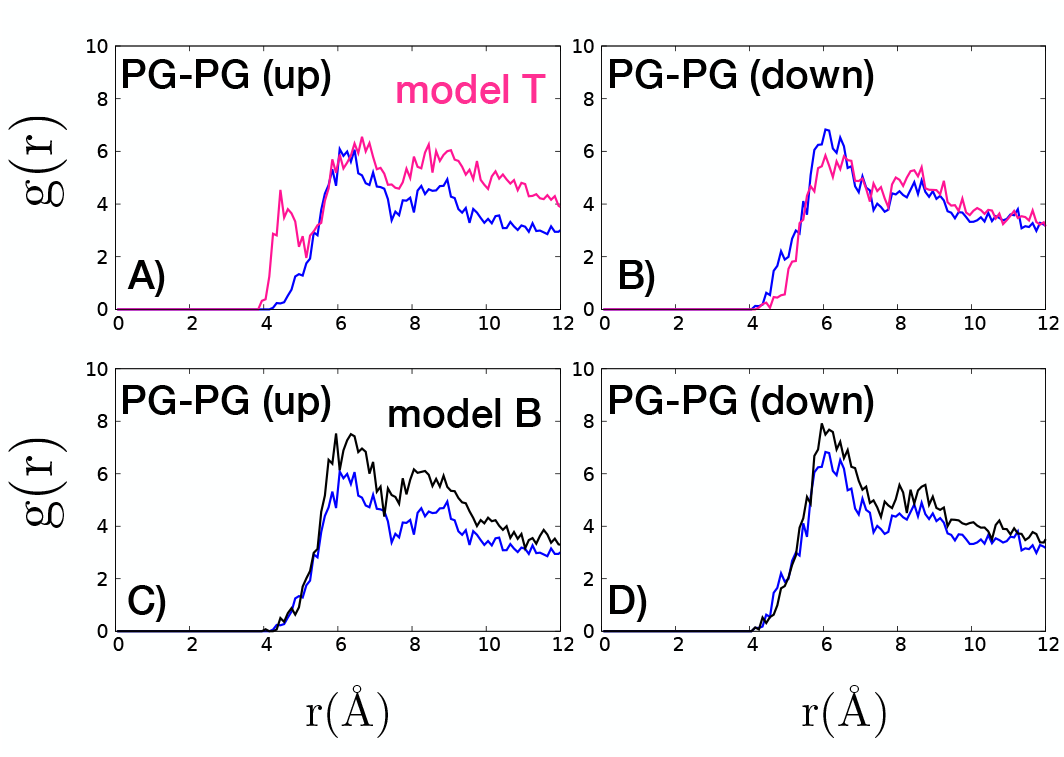
g(r) values measured between the phosphate groups of POPG-POPG lipids, averaged over 50 ns (650-700 ns) for the upper (denoted up) and lower leaflets (denoted down) of the model T and model B polymer-membrane systems. It is to be noted that the polymers interact with what is designated as upper leaflet. For comparison, g(r) for membrane only system is shown in blue color.

**Fig. 11.**
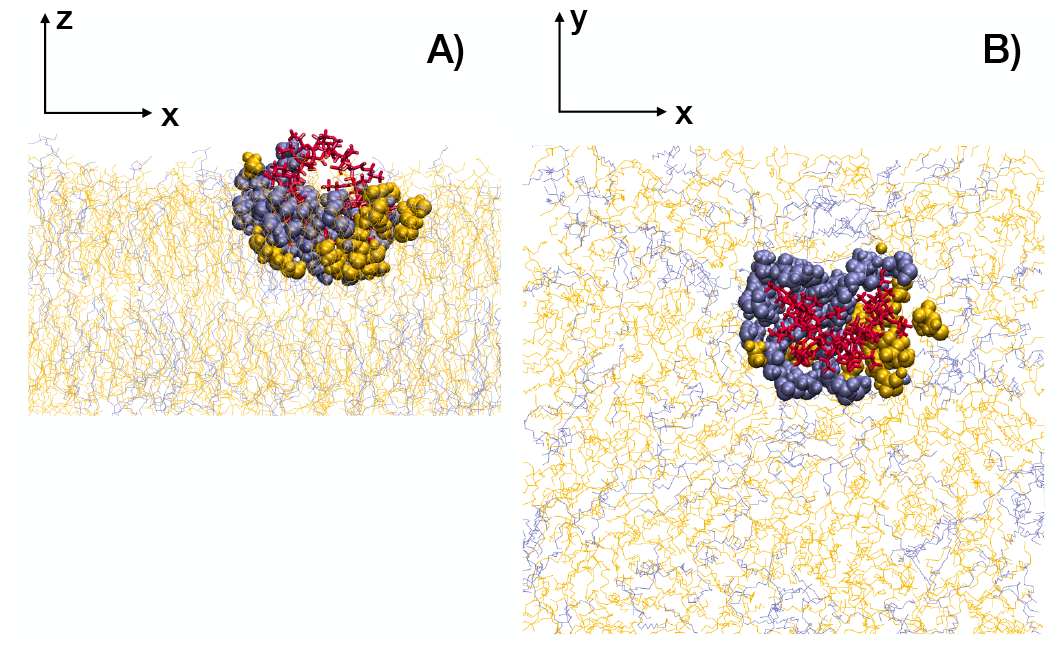
Snapshot of model T polymer inserted into the bacterial membrane at the end of 900 ns. The lipid molecules within 5 Å of the polymer are shown in vdW representation, rest of the lipids are in line representaion. POPE, POPG and polymer model T are colored orange, iceblue and red respectively. POPG seemingly displays higher local concentration around the ternary polymer.

A natural question now is to examine the change in the physical configuration of the membrane induced by the random ternary polymer. For this, we first investigate the lateral inhomogeinities in the membrane thickness induced by the model T polymer, which we compare with the control system (with no polymer interaction). For this, we plot the thickness profiles across the plane of the membrane, averaged over last 50 ns of simulations, as shown in Fig. 12. The 2-D thickness map is calculated using MEMBPLUGIN ^61^ extension in VMD by interpolating the distance between the phosphate groups of the upper and lower leaflet into the orthogonal grid in the X-Y plane of the bilayer with a grid spacing of 2Å. We observe that the thickness profile for the control system is largely uniform with average overall bilayer thickness around 38.37 ± 0.157 Å. On the other hand, in presence of the model T polymer, a clear upward shift in the average thickness of the bilayer is observed, with average bilayer thickness around 40.400 ± 0.162 Å. The bilayer thickness distribution also shows a considerable non-uniformity throughout the 2-D plane in the presence of ternary polymer, with local thickness value ranging from 35.5 to 42.5 Å, as shown in Fig. 12A. Such lateral inhomogeneity in membrane thickness has been shown to be related to the coarsening of the bilayer leaflet, leading to clustering of anionic and zwitterionic lipids which promote antimicrobial activities ^16,57^.

**Fig. 12.**
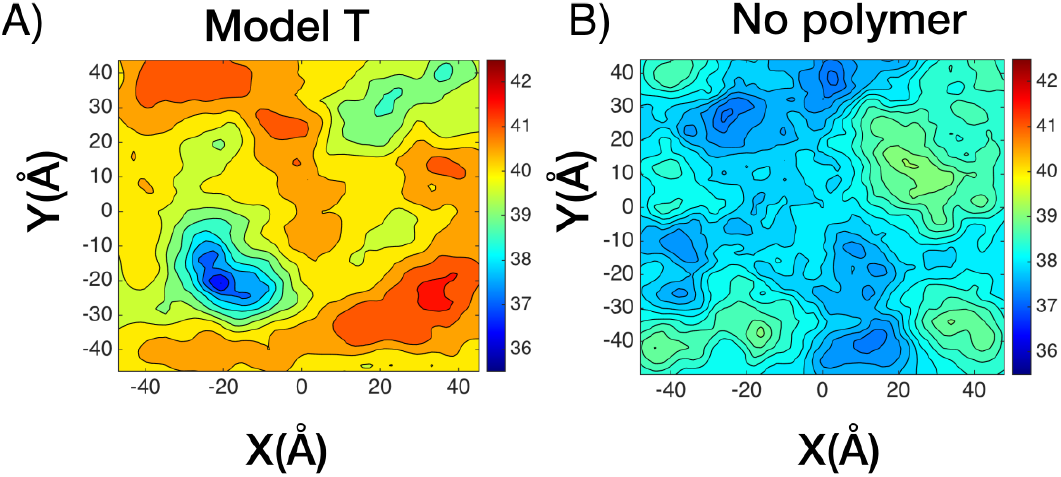
Contour plots showing the distributions of local membrane thickness in the X-Y plane for A) membrane in the presence of ternary polymer (model T) and B) only membrane (no polymer) system. Bilayer thickness is reported in Å.

Further, in previous work ^17^, it has been exhibited that the presence of multiple E4 polymers with aminobutylene cationic side chains in the bilayer reduces the average area per lipid. Strikingly this phenomenon is also observed in our case, with a single random ternary polymer, as well, with the area per lipid for the model T-membrane system (60.212 ± 0.076)Å^2^ being smaller as compared to the control (bilayer with no polymer) system (64.4 ± 0.108)Å^2^. Thus, our analysis shows that even a single random ternary polymer, upon membrane insertion, can induce extensive lipid reorganisation, which can then affect the physical configuration of the membrane.

### 3.5 Interaction of random polymer aggregates with bacterial membrane

In this section, we examine interactions of model T and model B polymer aggregates with the bacterial membrane. Both models T and B polymers form aggregates in solution phase, albeit with markedly different morphologies and inter-aggregate interactions ^62^. Illustrative configurations of the aggregates in solution, at the end of 150 ns of simulation runs, are shown in Fig. 13A and Fig. 13B. The formation and stability of such aggregates crucially depends on the hydrophobic content of the individual polymers. Strong aggregations are formed in case of random binary polymers, primarily driven by attractive interactions between hydrophobic groups. However, replacing some of the hydrophobic groups with overall charge neutral polar groups weakens the aggregate considerably, leading to increased conformational fluctuations and formation of loose-packed aggregates, in the case of random ternary polymers ^62^.

**Fig. 13.**
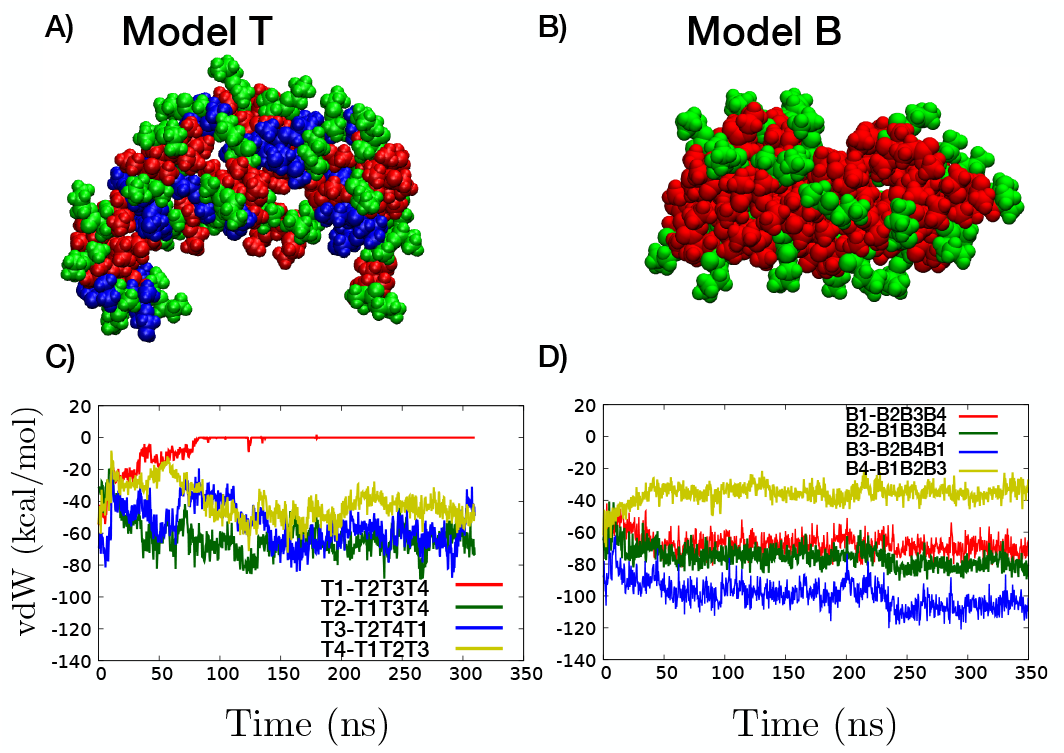
Representative configuration of an aggregate in solution phase, in case of A) ternary polymer model T and B) binary polymer model B. The cationic, hydrophobic and polar groups are colored green, red and blue respectively. Evolution of van der Waals interaction energy with simulation time between each polymer and all others, is plotted for C) model T and D) model B aggregates in lipid phase.

After placing the aggregates of model T and model B polymers over model bacterial membrane, we observe that both the aggregates approach the membrane surface within a few nanoseconds of simulation time, due to the attractive electrostatic interactions between the charged AEMA groups and the negatively charged head groups of POPG lipid molecules. However, subsequent membrane interaction is substantially different for the ternary and binary aggregates. We first probe the stability of the aggregates while interacting with the membrane by computing the van der Waals interaction energy as a function of simulation time (Fig. 13C and Fig. 13D), showing the interaction energy between each polymer with all other polymers in the aggregate. Clearly, model B polymers are decidedly stable in the aggregate and show robust attractive interactions among each other. In Fig. 14D, the final configuration of the model B aggregate at *t* = 350 ns of simulation time is shown, with the aggregate lying at the water-membrane interface with no polymer dissociation from aggregate and no partitioning into membrane interior. On the other hand, model T polymers show much weaker attraction among themselves with fluctuating interactions. We note that one of the polymer in the ternary aggregate, T1, shows continuous decrease in interaction with other polymers in the aggregate and eventually dissociates at ~ 85 ns from the aggregate (Fig. 14A). Around the same timescale, the polymer (T1)-lipid interactions become more favourable than the polymer (T1)-polymers (T2T3T4) interactions, as shown in Fig. 14B. It is also notable that the ternary polymer T1 shows identical membrane insertion mode as in the case of the single polymer simulations for the model T polymer (see Fig. 14C and Fig. 2B). Overall, this highlights the importance of the formation of “weak aggregates” ^62^, in aiding polymer dissociation and partitioning into bacterial membrane, thus strongly indicating the efficacy of ternary polymers as effective antimicrobial agents.

**Fig. 14.**
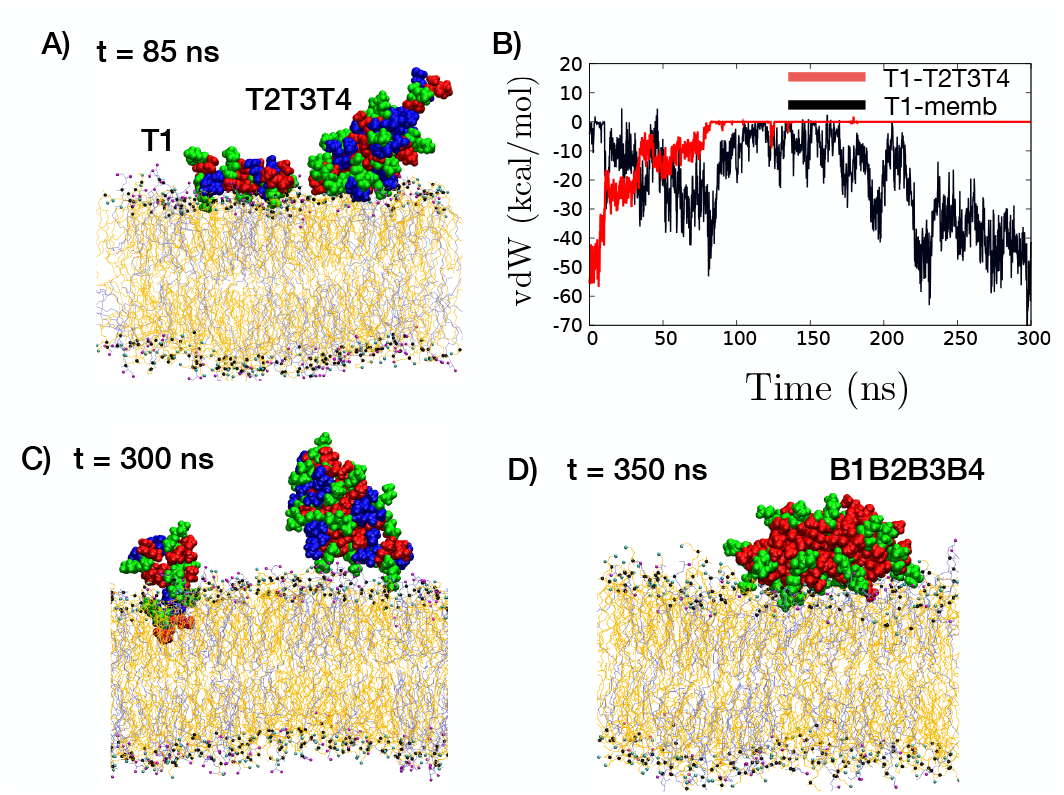
A) Representative snapshot showing separation of polymer T1 from rest of the three polymer for random ternary polymer aggregate in lipid phase. The separation happens in the timescale of ~ 85 ns. B) van der Waals interaction energy between polymer T1 and all other polymers in the ternary polymer aggregate (red color) and between polymer T1 and membrane lipids (black color). C) and D) illustrate final snapshots for ternary and binary aggregate at the end of simulation runs. The cationic, hydrophobic and polar groups are colored green, red and blue respectively. The POPE lipids are coloured orange and POPG lipids are shown in ice-blue colour; the lipid head group atoms are oxygen (magenta), nitrogen (cyan) and phosphate (black).

## 4 Discussion

In this report, we explored the interaction of ternary methacrylate biomimetic antimicrobial polymers, composed of hydrophobic, cationic and neutral polar functional groups, on model bacterial membrane. To understand the role played by the neutral polar groups in influencing the membrane interaction of such polymers, we compared it to the membrane interactions of traditional binary AM polymers, which are composed only of hydrophobic and cationic functional groups. Further, to probe the effect of sequence of the functional groups along the polymer chain, the ternary polymers were taken in two configurations, one in which the groups are randomly distributed along the backbone while other in which the polar groups were arranged in a block. Our results show that ternary polymers, specifically those in which the constituent functional groups are randomly distributed, can partition into the membrane, forming globular structure, while inducing significant lipid reorganisation in its vicinity and change the physical configuration of the membrane. It has been shown that localization of anionic PG lipid head groups induced by presence of antibacterial agents, which is followed by coarsening of the bilayer and consequent phase boundary defects, adversely affects the integrity of the bacterial membranes leading to cell lysis ^57–59^. Therefore, we can deduce that the presence of the ternary methacrylate antimicrobial polymer in membrane phase will likely result in disruption of membrane, fostering a robust antimicrobial response of such polymers.

It is also notable that the membrane insertion mode of the ternary polymer is markedly different from the binary polymer case. The binary polymer displays acquired amphiphilicity upon insertion into the bilayer, aligning parallel to the membrane surface with the cationic groups interacting with the lipid head groups and the hydrophobic groups interacting with the hydrophobic membrane core. On the other hand, in case of the ternary polymer, such a clear segregation of groups is absent, specifically in case of the random ternary polymer. In this case, the polymer stays aligned in the direction of the membrane normal as it penetrates the bilayer while adopting a folded conformation. This effect is underpinned by two competitive effects involving the presence of the neutral polar groups-attractive interactions with water on one hand and attractive interactions with membrane lipids on the other. Such an effect is however specific to the random ternary polymer, with a block arrangement of the polar groups resulting in a decidedly strong preference for interactions with water, which results in the block of polar groups largely lying in the aqueous environment above the membrane-water interface, impeding a deeper penetration of the polymer into the bilayer. We note here that in general the wide variety of AMPs have varying modes of membrane insertion ^63^, with for instance, some AMPs adopting a parallel alignment on the membrane surface like LL-37^64^ and indolicidin ^65^, while some other AMPs inserting into the bilayer perpendicular to the membrane surface, e.g. magainin 2 and lacticin Q ^66^. Such effects are likely a result of variations in the sequences of the AMPs due to the presence of different functional groups. Our results here also suggest that the classic conformation of facially amphiphilic structures is not prerequisite to effective attractive interactions between the antimicrobial agent and the bacterial membrane leading to possible new modes of antimicrobial mechanism.

Finally, we also studied the interaction of aggregates of binary and ternary polymers with model bacterial membrane. We exhibited that in case of the ternary polymers, their weak aggregation in solution results in fairly rapid disintegration of polymer from the aggregate, with the polymer subsequently penetrating the membrane, following a similar pattern as the membrane interactions of single ternary polymer. On the other hand, binary polymers, which form strong aggregates, failed to separate from the aggregate to interact with the membrane under similar conditions. It is notable that in previous work ^17^, interaction of aggregates of binary E4 polymers, having aminobutylene side chains, with bacterial membrane patch had been studied using MD simulations. It was shown in this case that polymers are released into the bilayer from the aggregate due to weak polymer-polymer interactions, which are overcome by polymer-anionic lipid interactions. However, these polymers have 7 charged cationic groups and 3 hydrophobic groups in a polymer chain (with degree of polymerization = 10) ^20^. The smaller hydrophobic content of the E4 polymers can explain the weaker aggregate formation, facilitating partitioning and membrane interactions. However, most natural AMPs have only 30% net positive charge ^40^ and experimental studies have also indicated that antimicrobial activity of polymers starts to level off when charged groups are more than ~ 30% of composition ^28^. Therefore, some charged groups in E4 polymer may be excessive and not necessarily required for potent antimicrobial activity. On the other hand, higher hydrophobic content clearly leads to formation of strong aggregates, as indicated by our results, obstructing efficient partitioning of the polymers and decreasing the efficacy of their antimicrobial action.

Therefore, in summary, our results indicate that ternary polymers, specifically those which have a random arrangement of functional groups, can effectively act as antibacterial agents due to their weak aggregation behaviour, which results in quick partitioning of polymers into the membrane, with these polymers then penetrating deep into the membrane while inducing significant lipid reorganisation in their vicinity as well as changes in membrane physical configuration. It is pertinent to note that natural AMPs typically have several more functional residues other than only charged, polar and hydrophobic units ^67–69^. The design of polymers, mimicking AMPs in displaying optimised compositions of a multitude of subunits, is of particular significance since it can lead to the development of potent antibacterial agents that can act against a broad spectrum of bacteria. Our present work fosters the design of such polymers.

## 5 Acknowledgments

All the simulations in this work have been carried out on clusters Annapurna and Nandadevi at The Institute of Mathematical Sciences, Chennai, India.

## References

1 H. W. Boucher, G. H. Talbot, J. S. Bradley, J. E. Edwards, D. Gilbert, L. B. Rice, M. Scheld, B. Spellberg and J. Bartlett, Clinical Infectious Diseases, 2009, 48, 1–12.

2 A.-P. Magiorakos, A. Srinivasan, R. Carey, Y. Carmeli, M. Falagas, C. Giske, S. Harbarth, J. Hindler, G. Kahlmeter, B. Olsson-Liljequist, D. Paterson, L. Rice, J. Stelling, M. Struelens, A. Vatopoulos, J. Weber and D. Monnet, Clinical Microbiology and Infection, 2012, 18, 268 – 281.

3 Y. Li, Q. Xiang, Q. Zhang, Y. Huang and Z. Su, Peptides, 2012, 37, 207–215.

4 A. Tossi, L. Sandri and A. Giangaspero, Peptide Science, 2000, 55, 4–30.

5 J. B. Mcphee and R. E. W. Hancock, Journal of Peptide Science, 2005, 11, 677–687.

6 R. E. W. Hancock and H.-G. Sahl, Nat Biotechnol, 2006, 24, 1551âĂŞ1557.

7 H. Khandelia and Y. N. Kaznessis, Biochimica et Biophysica Acta (BBA) - Biomembranes, 2007, 1768, 509 – 520.

8 J. Mondal, X. Zhu, Q. Cui and A. Yethiraj, The Journal of Physical Chemistry B, 2010, 114, 13585–13592.

9 J. N. Horn, T. D. Romo and A. Grossfield, Biochemistry, 2013, 52, 5604–5610.

10 C. M. Goodman, S. Choi, S. Shandler and W. F. DeGrado, Nature Chemical Biology, 2007, 3, 252–262.

11 G. N. Tew, R. W. Scott, M. L. Klein and W. F. DeGrado, Accounts of Chemical Research, 2010, 43, 30–39.

12 K. Kuroda and G. A. Caputo, WIREs Nanomedicine and Nanobiotechnology, 2013, 5, 49–66.

13 U. Baul and S. Vemparala, Antimicrobial Materials for Biomedical Applications, The Royal Society of Chemistry, 2019, pp. 113–136.

14 G. Rossi and L. Monticelli, Journal of physics. Condensed matter: an Institute of Physics journal, 2014, 26, 503101.

15 Y. Oda, K. Yasuhara, S. Kanaoka, T. Sato, S. Aoshima and K. Kuroda, Polymers, 2018, 10, 93.

16 J. N. Horn, J. D. Sengillo, D. Lin, T. D. Romo and A. Grossfield, Biochimica et Biophysica Acta (BBA) - Biomembranes, 2012, 1818, 212 – 218.

17 U. Baul, K. Kuroda and S. Vemparala, The Journal of Chemical Physics, 2014, 141, 084902.

18 C. Ergene, K. Yasuhara and E. F. Palermo, Polym. Chem., 2018, 9, 2407–2427.

19 K. D. Saint Jean, K. D. Henderson, C. L. Chrom, L. E. Abiuso, L. M. Renn and G. A. Caputo, Probiotics and antimicrobial proteins, 2018, 10, 408–419.

20 E. F. Palermo, S. Vemparala and K. Kuroda, Biomacromolecules, 2012, 13, 1632–1641.

21 Y. Yang, Z. Cai, Z. Huang, X. Tang and X. Zhang, Polymer Journal, 2018, 50, 33–44.

22 E. F. Palermo and K. Kuroda, Applied microbiology and biotechnology, 2010, 87, 1605–1615.

23 H. Takahashi, G. A. Caputo, S. Vemparala and K. Kuroda, Bioconjugate Chemistry, 2017, 28, 1340–1350.

24 D. S. S. M. Uppu, S. Samaddar, J. Hoque, M. M. Konai, P. Krishnamoorthy, B. R. Shome and J. Haldar, Biomacromolecules, 2016, 17, 3094–3102.

25 M. S. Ganewatta and C. Tang, Polymer (Korea), 2015, 63, A1–A29.

26 Y. Yang, Z. Cai, Z. Huang, X. Tang and X. Zhang, Polymer Journals, 2018, 50, 33–44.

27 K. Hu, N. W. Schmidt, R. Zhu, Y. Jiang, G. H. Lai, G. Wei, E. F. Palermo, K. Kuroda, G. C. L. Wong and L. Yang, Macromolecules, 2013, 46, 1908–1915.

28 H. Mortazavian, L. L. Foster, R. Bhat, S. Patel and K. Kuroda, Biomacromolecules, 2018, 19, 4370–4378.

29 B. C. Allison, B. M. Applegate and J. P. Youngblood, Biomacromolecules, 2007, 8, 2995–2999.

30 G. Wang and B. Mishra, Frontiers in immunology, 2012, 3, 221.

31 X. Yang, K. Hu, G. Hu, D. Shi, Y. Jiang, L. Hui, R. Zhu, Y. Xie and L. Yang, Biomacromolecules, 2014, 15, 3267–3277.

32 S. Chakraborty, R. Liu, Z. Hayouka, X. Chen, J. Ehrhardt, Q. Lu, E. Burke, Y. Yang, B. Weisblum, G. C. L. Wong, K. S. Masters and S. H. Gellman, Journal of the American Chemical Society, 2014, 136, 14530–14535.

33 K. Wakamatsu, A. Takeda, T. Tachi and K. Matsuzaki, Biopolymers, 2002, 64, 314–327.

34 K. Kuroda and W. F. DeGrado, Journal of the American Chemical Society, 2005, 127, 4128–4129.

35 T. Kouno, N. Fujitani, M. Mizuguchi, T. Osaki, S.-i. Nishimura, S.-i. Kawabata, T. Aizawa, M. Demura, K. Nitta and K. Kawano, Biochemistry, 2008, 47, 10611–10619.

36 D. Takahashi, S. K. Shukla, O. Prakash and G. Zhang, Biochimie, 2010, 92, 1236 – 1241.

37 C. L. Chrom, L. M. Renn and G. A. Caputo, Antibiotics, 2019, 8, 20.

38 G. N. Tew, D. Liu, B. Chen, R. J. Doerksen, J. Kaplan, P. J. Carroll, M. L. Klein and W. F. DeGrado, Proceedings of the National Academy of Sciences, 2002, 99, 5110–5114.

39 M. A. Gelman, B. Weisblum, D. M. Lynn and S. H. Gellman, Organic Letters, 2004, 6, 557–560.

40 M. Zasloff, Nature, 2002, 415, 389–395.

41 A. A. Polyansky, R. Ramaswamy, P. E. Volynsky, I. F. Sbalzarini, S. J. Marrink and R. G. Efremov, The Journal of Physical Chemistry Letters, 2010, 1, 3108–3111.

42 S. Jo, J. B. Lim, J. B. Klauda and W. Im, Biophysical Journal, 2009, 97, 50 – 58.

43 U. Baul and S. Vemparala, Soft Matter, 2017, 13, 7665–7676.

44 W. L. Jorgensen, J. Chandrasekhar, J. D. Madura, R. W. Impey and M. L. Klein, The Journal of Chemical Physics, 1983, 79, 926–935.

45 J. C. Phillips, R. Braun, W. Wang, J. Gumbart, E. Tajkhor-shid, E. Villa, C. Chipot, R. D. Skeel, L. Kale and K. Schulten, Journal of Computational Chemistry, 2005, 26, 1781 – 1802.

46 J. B. Klauda, R. M. Venable, J. A. Freites, J. W. O’ Connor, D. J. Tobias, C. Mondragon-Ramirez, I. Vorobyov, A. D. MacKerell and R. W. Pastor, The Journal of Physical Chemistry B, 2010, 114, 7830–7843.

47 A. D. MacKerell, D. Bashford, M. Bellott, R. L. Dunbrack, J. D. Evanseck, M. J. Field, S. Fischer, J. Gao, H. Guo, S. Ha, D. Joseph-McCarthy, L. Kuchnir, K. Kuczera, F. T. K. Lau, C. Mattos, S. Michnick, T. Ngo, D. T. Nguyen, B. Prodhom, W. E. Reiher, B. Roux, M. Schlenkrich, J. C. Smith, R. Stote, J. Straub, M. Watanabe, J. WiÃşrkiewicz-Kuczera, D. Yin and M. Karplus, The Journal of Physical Chemistry B, 1998, 102, 3586–3616.

48 I. Ivanov, S. Vemparala, V. Pophristic, K. Kuroda, W. F. De-Grado, J. A. McCammon and M. L. Klein, Journal of the American Chemical Society, 2006, 128, 1778–1779.

49 H. Seeger, G. Marino, A. Alessandrini and P. Facci, Biophysical Journal, 2009, 97, 1067 – 1076.

50 G. J. Martyna, D. J. Tobias and M. L. Klein, The Journal of Chemical Physics, 1994, 101, 4177–4189.

51 S. E. Feller, Y. Zhang, R. W. Pastor and B. R. Brooks, The Journal of Chemical Physics, 1995, 103, 4613–4621.

52 U. Essmann, L. Perera, M. L. Berkowitz, T. Darden, H. Lee and L. G. Pedersen, The Journal of Chemical Physics, 1995, 103, 8577–8593.

53 W. Humphrey, A. Dalke and K. Schulten, Journal of Molecular Graphics, 1996, 14, 33 – 38.

54 E. F. Palermo, S. Vemparala and K. Kuroda, in Antimicrobial Polymers: Molecular Design as Synthetic Mimics of Host-Defense Peptides, American Chemical Society, 2013, ch. 20, pp. 319–330.

55 M. A. Rahman, M. Bam, E. Luat, M. S. Jui, M. S. Ganewatta, T. Shokfai, M. Nagarkatti, A. W. Decho and C. Tang, Nature Communications, 2018, 9, 5231.

56 F. Jean-François, S. Castano, B. Desbat, B. Odaert, M. Roux, M.-H. Metz-Boutigue and E. J. Dufourc, Biochemistry, 2008, 47, 6394–6402.

57 R. F. Epand, B. P. Mowery, S. E. Lee, S. S. Stahl, R. I. Lehrer, S. H. Gellman and R. M. Epand, Journal of Molecular Biology, 2008, 379, 38 – 50.

58 R. M. Epand and R. F. Epand, Biochimica et Biophysica Acta (BBA) - Biomembranes, 2009, 1788, 289 – 294.

59 P. Joanne, C. Galanth, N. Goasdoué, P. Nicolas, S. Sagan, S. Lavielle, G. Chassaing, C. El Amri and I. D. Alves, Biochimica et Biophysica Acta (BBA) - Biomembranes, 2009, 1788, 1772 – 1781.

60 W. Zhao, T. Róg, A. Gurtovenko, I. Vattulainen and M. Karttunen, Biochimie, 2008, 90, 930–8.

61 Guixà-González, Ramon and Rodriguez-Espigares, Ismael and Ramírez-Anguita, Juan Manuel and Carrió-Gaspar, Pau and Martinez-Seara, Hector and Giorgino, Toni and Selent, Jana, Bioinformatics, 2014, 30, 1478–1480.

62 G. Rani, K. Kuroda and S. Vemparala, Journal of Physics: Condensed Matter (accepted for publication), 2020.

63 P. Kumar, J. Kizhakkedathu and S. Straus, Biomolecules, 2018, 8, 4.

64 Y. Shai, Peptide Science, 2002, 66, 236–248.

65 A. Rozek, C. Friedrich and R. Hancock, Biochemistry, 2001, 39, 15765–74.

66 T.-H. Lee, K. Hall and M.-I. Aguilar, Current topics in medicinal chemistry, 2015, 16,.

67 H. Sato and J. B. Feix, Biochimica et Biophysica Acta (BBA) - Biomembranes, 2006, 1758, 1245 – 1256.

68 S. Colak, C. F. Nelson, K. Nüsslein and G. N. Tew, Biomacromolecules, 2009, 10, 353–359.

69 E. H. H. Wong, M. M. Khin, V. Ravikumar, Z. Si, S. A. Rice and M. B. Chan-Park, Biomacromolecules, 2016, 17, 1170–1178.

